# *Bacillus subtilis* encodes a discrete flap endonuclease that cleaves RNA-DNA hybrids

**DOI:** 10.1101/2022.12.20.521345

**Authors:** Frances Caroline Lowder, Lyle A. Simmons

## Abstract

Current models for Okazaki fragment maturation in eubacteria invoke RNA cleavage by RNase H, followed by strand displacement synthesis and 5′ RNA flap removal by DNA polymerase I (Pol I). RNA removal by Pol I is thought to occur through the 5′-3′ flap <u>e</u>ndo/exo<u>n</u>uclease (FEN) domain, located in the N-terminus of the protein. In addition to Pol I, many bacteria encode a second, Pol I-independent FEN. The contribution of Pol I and Pol I-independent FENs to DNA replication and genome stability remains unclear. In this work we purified Pol I and FEN, then assayed these proteins on a variety of RNA-DNA hybrid and DNA-only substrates. We found that FEN is far more active than Pol I on 5′ flapped and nicked RNA-DNA hybrid substrates. We found that the 5′ nuclease activity of *B. subtilis* Pol I is feeble, even during DNA synthesis when a 5′ flapped substrate is formed modeling an Okazaki fragment intermediate. Examination of Pol I and FEN on DNA-only substrates shows that FEN is more active than Pol I on most substrates tested. Further experiments show that Δ*polA* phenotypes are completely rescued by expressing the C-terminal polymerase domain while expression of the N-terminal 5′ nuclease domain fails to complement Δ*polA*. Cells lacking FEN (Δ*fenA*) show a phenotype in conjunction with an RNase HIII defect, providing genetic evidence for the involvement of FEN in Okazaki fragment processing. With these results, we propose a model where cells remove RNA primers using FEN while upstream Okazaki fragments are extended through synthesis by Pol I. Our model is similar to models for Okazaki fragment processing in eukaryotes, where Pol d catalyzes strand displacement synthesis followed by 5′ flap cleavage using FEN-1.

**Author Summary:** 5′ flap endo/exonuclease (FEN) activity provides an essential contribution to DNA replication and repair in all cellular life. In bacteria, DNA polymerase I is thought to be the central enzyme involved in Okazaki fragment processing, using its DNA polymerase and 5′ nuclease activities to generate and then remove the 5’ ssRNA segment of an Okazaki fragment. Many bacterial genomes encode a second, discrete FEN in addition to Pol I. We show that FEN is the primary 5′ nuclease used by *B. subtilis* for primer removal. FEN activity exceeds that of Pol I on most substrates, including several that mimic Okazaki fragment intermediates. Additionally, we provide genetic evidence showing that FEN is involved in Okazaki fragment processing and that it is the DNA polymerase domain of Pol I rather than its 5′ nuclease domain that is important *in vivo*. With our results, we propose a new model for Okazaki fragment processing in *B. subtilis*, which may be prevalent in a wider group of bacteria.

## Introduction

RNA-DNA hybrids form during transcription and DNA replication in all cells and many viruses. While their formation is essential, persistence of RNA-DNA hybrids can lead to genome instability by causing nicks and double stranded breaks when the 2′ OH on the ribose sugar reacts with the phosphodiester bond, forming a 2′, 3′ cyclic phosphate [1]. During normal cell growth, RNA-DNA hybrids occur in several different forms within cells. R-loops represent one type of RNA-DNA hybrid, occurring when newly transcribed mRNA remains base-paired with the complementary DNA strand following transcription [2,3]. RNA-DNA hybrids also occur when ribonucleotides are covalently joined to DNA. Sugar errors can occur when single ribonucleotide monophosphates (rNMPs) or stretches of rNMPs are covalently nested in DNA by replicative polymerases [4,5] or under stress conditions when error-prone synthesis occurs [6]. Covalent RNA-DNA hybrids also form when primase synthesizes RNA primers at the initiation of leading strand synthesis and for synthesis of each Okazaki fragment during lagging strand replication [7–9]. Due to the frequency of Okazaki fragment synthesis, RNA primers occur predominantly on the lagging strand, resulting in the incorporation of approximately 20,000 rNMPs per genome replication event [10]. Because RNA-DNA hybrids can lead to genomic instability and disease, all cells enlist a wide array of proteins that recognize and resolve the different types of hybrids that form *in vivo* [11]. The ribonuclease H (RNase H) proteins are a well characterized group of enzymes involved in the repair of all types of RNA-DNA hybrids that occur in cells. RNase HI, HII, and HIII are all capable of cleaving substrates with stretches of four or more ribonucleotides, which includes the RNA in Okazaki fragments. RNase HI and HIII are also able to resolve R-loops, while RNase HII is responsible for cleaving at single ribonucleotide errors in genomic DNA [12,13].

Several studies have provided evidence that RNase Hs participate in the removal of RNA primers during Okazaki fragment maturation [12–14], however, studies in *E. coli* identified DNA polymerase I (Pol I) as the major bacterial protein involved in this process [14,15]. Pol I is composed of three domains: a 5′-3′ polymerase, a 3′-5′ exonuclease, and a 5′-3′ nuclease [16,17]. The well-studied Klenow fragment consists of the 5′-3′ polymerase and 3′-5′ exonuclease located in the C-terminus of the protein and is responsible for DNA synthesis and error “proofreading”, while the 5′-3′ nuclease domain is located at the N-terminus of Pol I [15–20]. Coordination of Pol I activities during lagging strand synthesis should allow Pol I to synthesize DNA from an upstream Okazaki fragment through the downstream RNA primer and subsequently remove the RNA primer. The remaining nick between adjacent DNA fragments would then be sealed by DNA ligase to complete the repair [15,21]. Due to the importance of Pol I activities during DNA replication, Pol I is very well conserved across bacterial species [22], although many bacteria, including *B. subtilis*, lack the catalytic residues associated with an active 3′-5′ exonuclease [23], suggesting that the polymerase activity and 5′ nuclease activity of Pol I are most critical *in vivo* [24]. Current models propose that Okazaki fragment maturation in eubacteria is primarily accomplished by Pol I [15]. This model stems from studies in *E. coli* and *B. subtilis* showing the accumulation of ribonucleotides in short DNA fragments from cells expressing various mutants of *polA* [25–28]. Based on these early studies, the overarching model for Okazaki fragment maturation in bacteria invokes RNA incision by RNase HI (or HIII) with DNA synthesis and the bulk of RNA removal catalyzed by Pol I.

More recent studies have used sequence and structural homology to identify the N-terminal nuclease domain of Pol I as part of a larger superfamily of proteins with 5′-3′ flap <u>e</u>ndo<u>n</u>uclease and 5′-3′ exonuclease activity, known as FENs [29]. FEN activity is essential for replication; as such, proteins in this family are found across all domains of life [30–37]. Notable features of the FEN family include the binding of two or three bivalent cations in the active site, a helical or unstructured arch near the active site, and a C-terminal DNA binding motif [30,38–41]. FENs are structure-specific nucleases best known for their endonuclease activity on 5′ flap overhang substrates generated during strand-displacement synthesis [42–45]. Proteins in the FEN family have also been shown to have activity on a variety of other substrates, including nicked duplex DNA [29,32,43]. In all three kingdoms of life, organisms exist that encode multiple FENs [29,37,46]. In many bacteria, one of the FENs is a part of Pol I while the other is an independently encoded protein [22,29,35]. The FENs found as a domain of Pol I represent the most well-studied bacterial FENs to date, while the discrete FENs have received much less attention, making their contribution to DNA replication and genome instability unclear.

*B. subtilis* is one such organism known to encode a discrete FEN in addition to Pol I. The protein YpcP was initially identified as a protein homologous to the N-terminal domain of Pol I [22,47]. While YpcP has been described as an exonuclease, due to its similarity to the N-terminal domain of Pol I [48], and renamed ExoR, sequence homology and structural predictions of the Pol I N-terminal domain and YpcP suggest that both proteins belong to the wider FEN family [29]. For this reason, we will adopt the original nomenclature proposed for Pol I-independent, bacterial FENs [29] and herein refer to *B. subtilis* YpcP as FEN and its gene as *fenA*. Initial work characterizing the relationship between Pol I and FEN demonstrated that *B. subtilis polA* and *fenA* mutants are synthetically lethal [35,47] and that overexpression of *polA* or *fenA* was able to partially suppress the filamentous phenotype of an RNase H-deficient strain [35]. However, overexpression of *fenA* was unable to rescue a temperature sensitive strain lacking the N-terminal domain of Pol I [35]. Through these experiments, it was concluded that the two proteins have overlapping functions but are not redundant. Biochemical assays using purified FEN suggest that it has nucleolytic activity on a variety of substrates, with some preference shown for RNA-DNA hybrids [13]. Other evidence from protein-DNA binding assays suggests that FEN has the highest binding affinity for double stranded DNA with a 5′ overhang, suggesting that FEN might prefer to act on DNA [48]. Together these data suggest that FEN may actively contribute to Okazaki fragment processing and DNA repair, although the relationship between Pol I and FEN remains unknown.

In this work, we examined the substrate preference for Pol I and FEN using a variety of RNA-DNA hybrids and DNA only substrates. We also examined the phenotype of *polA* and *fenA* during genotoxic stress and in RNase H deficient backgrounds to determine the contribution of Pol I and FEN 5′ nuclease activity to genome integrity. We found that FEN shows the most robust activity on RNA-DNA hybrid substrates modeling Okazaki fragment intermediates and that the strong *polA* phenotypes are rescued by simply expressing the C-terminal domain lacking 5′ nuclease activity. With our results, we conclude that discrete *B. subtilis* FEN functions as the major Okazaki fragment nuclease, with Pol I DNA polymerase activity contributing to both DNA repair and Okazaki fragment resynthesis. Because discrete FENs are present in a wide-range of bacterial genomes, we suggest that Pol I-independent FENs provide the 5′ nuclease activity important for RNA removal during lagging strand replication in many bacteria. Our work suggests that lagging strand processing in these bacteria occurs through a process more similar to eukaryotes than previously appreciated.

## Results

### Loss of *fenA* confers sensitivity to DNA damage in the absence of ribonuclease HIII

Since *B. subtilis* encodes two proteins with hypothesized FEN activity, we began by investigating the importance of *fenA* for growth during conditions that result in DNA damage or dNTP depletion to test the model that FEN contributes to genome maintenance under stress conditions [48,49]. As shown in **Fig 1A**, a single deletion of *fenA* does not result in any detectable phenotype for cells grown on hydroxyurea (HU) or cells exposed to UV radiation (**S1 Fig**). It was previously shown that *fenA* disruption does not confer cold sensitivity nor sensitivity to mitomycin C (MMC) or methyl methanesulfonate (MMS) [13]. Given these results, it seems clear that *B. subtilis* FEN does not have an appreciable role during DNA repair *in vivo*. Another model is that FEN is involved in Okazaki fragment maturation. If true, we would expect a strain lacking both *rnhC* and *fenA* to confer a phenotype. Ribonuclease HIII (RNHIII), encoded by *rnhC*, is a protein known to participate in Okazaki fragment maturation and the resolution of RNA-DNA hybrids [12,13]. The absence of both proteins resulted in cells that were more sensitive to HU or UV than either single deletion (**Fig 1A and S1 Fig**), suggesting that FEN could be involved in the resolution of RNA-DNA hybrids on the lagging strand, and any defects due to the absence of FEN are exacerbated with the combined absence of RNHIII and presence of genotoxic stress.

**Fig 1.**
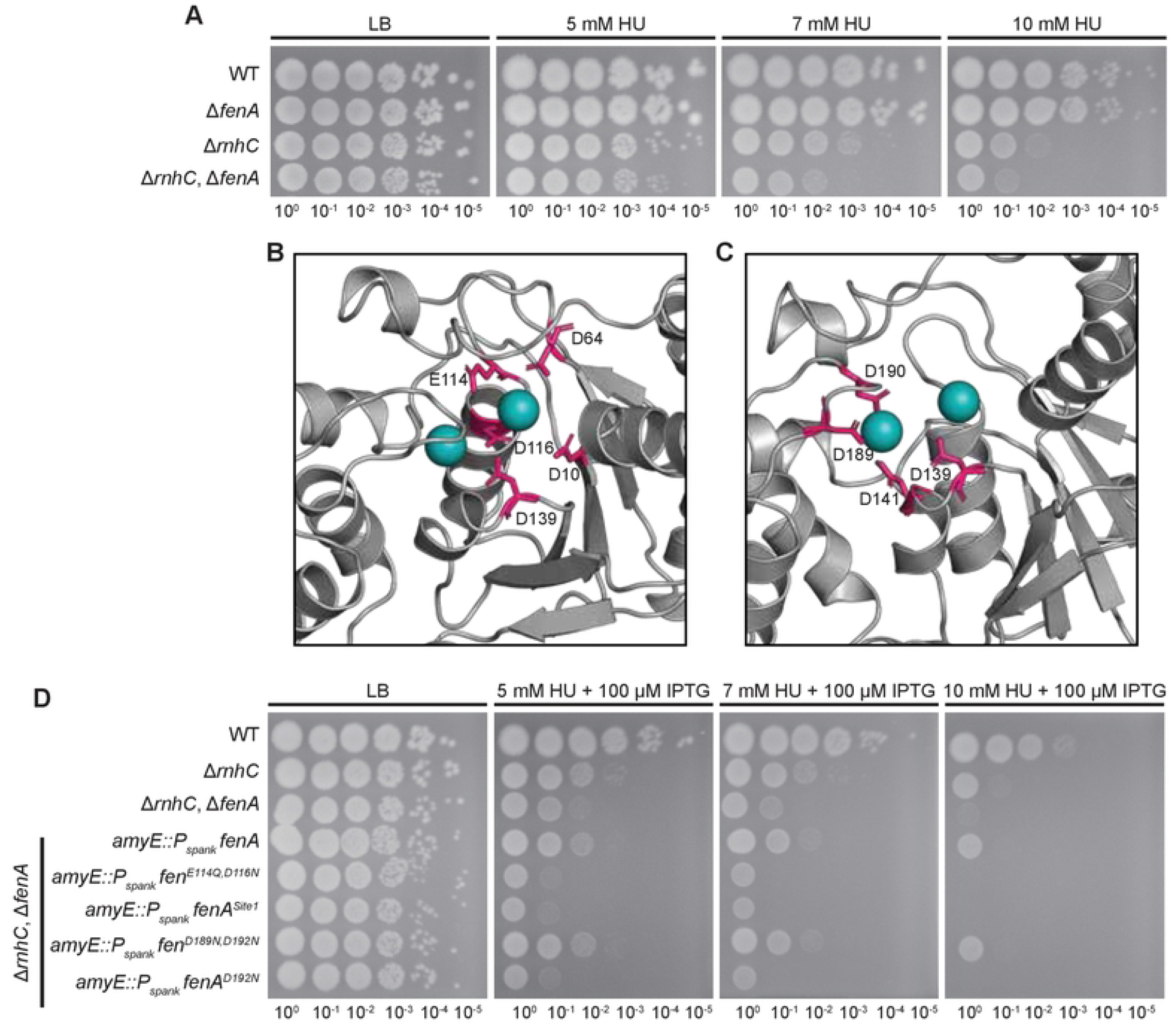
FEN contributes to RNA-DNA hybrid repair *in vivo*. (A) Spot titer assay to determine a phenotype of cells exposed to hydroxyurea in the absence of *fenA. rnhC* encodes Ribonuclease HIII, a protein known to be involved in the resolution of RNA-DNA hybrids. (B) Alphafold model of FEN, with Site 1 conserved carboxylate residues and (C) Site 2 carboxylate residues shown in pink. Coordinated bivalent metals are shown in blue and were added via alignment to *Mycobacterium smegmatis* FEN (PDB: 6C33). (D) Spot titer assay testing ectopically expressed mutants of *fenA* lacking conserved carboxylate residues for rescue of Δ*rnhC*, Δ*fenA* HU sensitivity.

### FEN is reliant on residues conserved across bacterial FENs

FEN contains a conserved group of eight carboxylate resides that are predicted to be a part of the active site [29], which is modeled in **Figs 1B and 1C**, with a multiple sequence alignment provided (**S2 Fig**). To test the importance of these amino acids, we designed a series of mutations to disrupt conserved residues and expressed each variant from the ectopic chromosomal *amyE* locus under the *Pspank* promoter, which allows induction of expression in the presence of isopropyl ß-D-1-thiogalactopyranoside (IPTG). As discussed above, the single deletion of *fenA* did not result in an obvious phenotype, so mutants were expressed in the Δ*rnhC*, Δ*fenA* background.

Overexpression of WT *fenA* rescued the double deletion phenotype (**Fig 1D and S3 Fig**). With the Δ*rnhC*, Δ*fenA* complementation assay established, four *fenA* mutants were created, two for each metal-binding site. The first mutant, *fenA*^E114Q,D116N^, was designed to abrogate binding of the Site 1 bivalent cation without changing the overall shape of the active site. The binding at Site 1 was also altered in the second mutant, where the EADD motif (residues 114 to 117) was changed to AAAA, a construct designated as *fenA*^Site1^. In addition to loss of cation binding, we predict that this mutant would have alterations to the shape of the active site due to the hydrophobic residues, as well as changes to the flexible arch, since sequence similarity suggests that the D117 residue interacts with R82 to form a salt-bridge [50]. As shown in **Fig 1D and S3 Fig**, neither mutant was able to rescue Δ*rnhC*, Δ*fenA* as well as WT *fenA*, suggesting that these residues are important for the function of FEN *in vivo*. The third mutant, *fenA*^D192N^, has one residue from metal-binding Site 2 changed. Previous work suggested that this mutant was catalytically inactive [13], however, overexpression in the double deletion strain rescues to nearly the same degree as the WT gene (**Fig 1D and S3 Fig**). Adding a second mutation, creating *fenA*^D189N,D192N^, once again led to a failure to rescue the Δ*rnhC*, Δ*fenA* strain. Together this suggests that not every conserved residue is critical for activity, however, perturbations that significantly affect metal-binding render *fenA* inactive *in vivo*.

An interesting observation from our experiments is that expression of *fenA*^E114Q,D116N^ or *fenA*^D189N,D192N^ in the Δ*rnhC*, Δ*fenA* strain results in cells that are more sensitive to HU than the parent strain or those expressing WT *fenA* (**Fig 1D**). To test if these mutations were dominant negative, we used the same system to express each mutant or the WT allele from an ectopic locus in WT or a Δ*rnhC* strain. There was no detectable phenotype in WT cells (**S4 Fig**), however, induced expression of either *fenA*^E114Q,D116N^ or *fenA*^D189N,D192N^ in the Δ*rnhC* background resulted in cells that were demonstrably more sensitive to HU than the parent strain (**S5 Fig**). Even when *fenA*^E114Q,D116N^ or *fenA*^D189N,D192N^ were induced on LB plates with no stressor, the colonies were smaller than those of the parent strain (**S5 Fig**). We speculate that the metal-binding variants are still able to bind substrates *in vivo* but are unable to bind one of the metal ions and therefore lack the catalytic activity required for turnover. We hypothesize that in WT cells, the mutants are outcompeted by the overlapping functions of Pol I, FEN, and RNHIII, and cause no phenotype. Loss of RNHIII disrupts this balance, which is then exacerbated by exposure to HU [51], ultimately leading to a phenotype during mutant overexpression.

### FEN is more active on 5′ flap substrates than Pol I

The *fenA* phenotype is most supportive of a contribution to Okazaki fragment processing as opposed to DNA repair. Given this, we chose to assay FEN and Pol I *in vitro* to compare their nuclease activity on a series of substrates to identify the substrates preferred by each enzyme. To do so, we expressed and purified each protein with a His_6_-SUMO tag, which we cleaved prior to use (detailed in Materials and Methods). This allowed purification of full-length proteins without any additional residues (**S6 Fig**). Using fluorescently labeled oligonucleotides (**S1 Table**), we generated a series of biologically relevant substrates. Each substrate had a DNA-only version, as well as a chimera / RNA-DNA hybrid version, where the first 12 bases at the 5′ end of the labeled strand are equivalent ribonucleoside monophoshates (rNMPs). In **Figs 2-7**, the assayed substrates are depicted at the top of the gels, with ribonucleotides indicated by zig-zag lines. As the activity of FENs is dependent on the bivalent cations at the protein’s active site, we used a buffer containing physiologically relevant concentrations of Mg^2+^ and Mn^2+^ as described previously [12].

**Fig 2.**
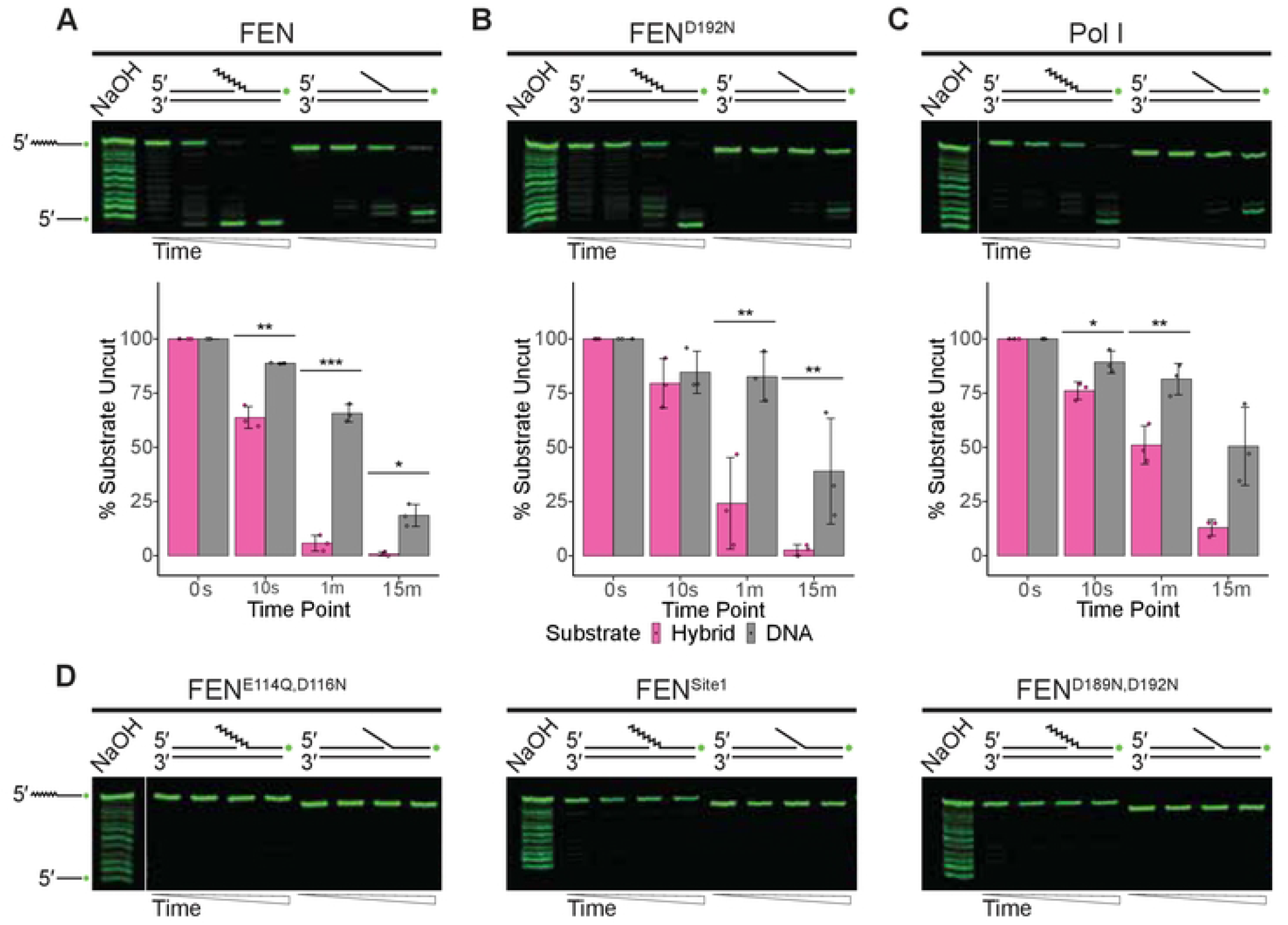
FEN is active on flap overhang structures. (A) Endonuclease activity of FEN, (B) FEN^D192N^, or (C) Pol I on 5′ flap overhang structures visualized using denaturing urea-PAGE. Percentage of substrate remaining uncut by each protein across three replicates is quantified with standard error bars beneath a representative gel; asterisks indicate significance: * p<0.05, ** p<0.01, or *** p<0.001. (D) Assays of activity on 5′ flap structures by the catalytic mutants FEN^E114Q,D116N^, FEN^Site1^, or FEN^D189N,D192N^. For all assays, flap substrates consist of oligonucleotides oJR366 and oJR368 with oJR339 (RNA-DNA hybrid; RNA indicated by zigzags) or oJR348 (DNA-only). Time points for all assays are 0 s, 10 s, 1 m, and 15 m with ladders generated by alkaline hydrolysis.

The first substrate we tested was a 5′ flap overhang, which consists of a double stranded substrate with a displaced, single-stranded 5′ arm (**Fig 2**). This is the canonical structure associated with FEN activity [52] and can be generated *in vivo* during strand-displacement synthesis by the polymerase domain of Pol I [53]. This structure is often associated with Okazaki-fragment maturation [27,28], nucleotide excision repair (NER) or base excision repair (BER) pathways [54]. As shown in **Fig 2A**, FEN was able to cleave both versions of this substrate well, rapidly generating a low molecular weight product for both, although it had significantly more activity on the hybrid flap structure. The FEN^D192N^ mutant, previously suggested to be catalytically inactive [13], had reduced activity on both substrates, however it generated similar products to that of the WT FEN (**Fig 2B**). Like FEN, Pol I preferred the hybrid flap substrate and generated a product consistent with flap endonuclease activity, however it had overall lower catalytic activity than FEN (**Fig 2C**). The three remaining mutants, FEN^E114Q,D116N^, FEN^Site1^, and FEN^D189N,D192N^ show no noticeable activity on the substrate, consistent with their failure to rescue Δ*rnhC*, Δ*fenA* strains (**Fig 1D**).

### FEN is more active on nicked substrates than Pol I

We next investigated nuclease activity on a nicked substrate (**Fig 3**). Nicks are a common type of DNA damage, where a missing phosphodiester bond occurs in otherwise continuous DNA. Nicks can result from many DNA repair pathways, including NER, BER, and mismatch repair [55]. Nicks can also be created following RNase H activity [56] or during Okazaki fragment maturation [57]. FEN showed high activity on the RNA-DNA hybrid nick substrate (**Fig 3A**), fully removing the rNMPs. While FEN was also fully active on the DNA-only substrate, it appears to use primarily 5′-3′ exonuclease activity, resulting in a sequential removal of nucleotides from the 5′ end rather than the single band associated with flap endonuclease activity. Regardless of the specific type of incision used, FEN had an overall preference for the hybrid structure. The FEN^D192N^ mutant (**Fig 3B**) shows both flap endonuclease and 5′ exonuclease activities on the hybrid structure, but not at the same rate observed for the WT enzyme. Despite the differences in rate, FEN and the FEN^D192N^ mutant generate similar products for each substrate by the 15-minute time point. Conversely, Pol I had markedly less activity on either variant of the nicked substrate than FEN (**Fig 3C**). Pol I engaged minimally in 5′ exonuclease activity on each nick, removing single nucleotides from each substrate. Interestingly, Pol I had significantly more activity on the DNA-only substrate than the hybrid, demonstrating a preference opposite that of FEN. As before, the remaining FEN mutants appear catalytically inactive and do not have detectable activity on the nicked substrate (**Fig 3D**).

**Fig 3.**
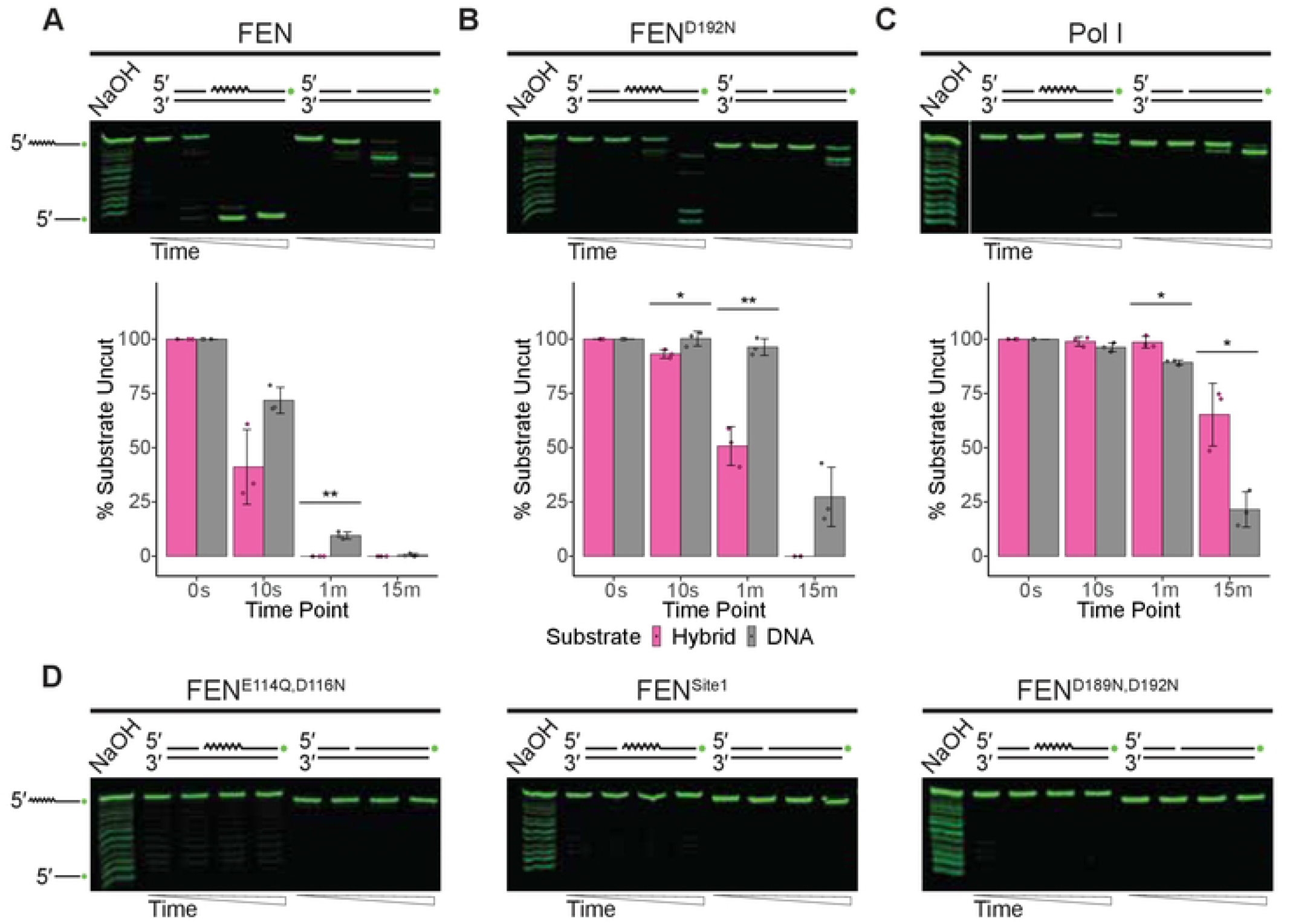
FEN can remove RNA from Okazaki fragment-like nicks. (A) Visualization of nuclease activity of FEN, (B) FEN^D192N^, or (C) Pol I on nicked duplex substrates. Three replicates of each assay are quantified beneath a representative urea-PAGE gel. Asterisks are used to indicate significance: * p<0.05, ** p<0.01, or *** p<0.001, while standard error bars are also provided. (D) Results of incubation of nicked duplex substrates with the FEN mutants FEN^E114Q,D116N^, FEN^Site1^, or FEN^D189N,D192N^. Nicked duplex substrates consist of oJR338 and oJR340 with oJR339 (RNA-DNA hybrid; RNA indicated by zigzags) or oJR348 (DNA). Ladders were generated by treating hybrid structure with sodium hydroxide. Assay time points for all reactions are 0 s, 10 s, 1 m, and 15 m.

### FEN is more active on 3′ overhang structures than Pol I

The third canonical substrate we tested was a 3′ overhang (**Fig 4**). This substrate is primarily associated with Okazaki fragment maturation [15] but can be formed during other cellular processes [58]. FEN was highly active on the hybrid version of this substrate, which most closely mimics an Okazaki fragment, cleaving all rNMPs from more than 80% of the substrate within 10 seconds (**Fig 4A**). FEN was also active on the DNA-only substrate, although the cleavage pattern suggests that this is primarily 5′ exonuclease activity. As with the other substrates, the FEN^D192N^ mutant had reduced activity on both the hybrid and DNA-only substrates, producing products consistent with those of WT FEN (**Fig 4B**). Pol I had minimal exonuclease activity on either substrate, engaging primarily in 5′ exonuclease activity (**Fig 4C**). Despite the high activity of WT FEN on the 3′ overhang, the remaining FEN mutants were not demonstrably active, further highlighting the essentiality of these residues to the protein’s function (**Fig 4D**). These substrates: 5′ flaps, nicked duplex DNA, and 3′ overhangs, represent the three most common structures associated with Okazaki fragment maturation. We conclude that the relative activities of FEN and Pol I on these structures suggest that FEN may contribute significantly to the resolution of these Okazaki fragments *in vivo*.

**Fig 4.**
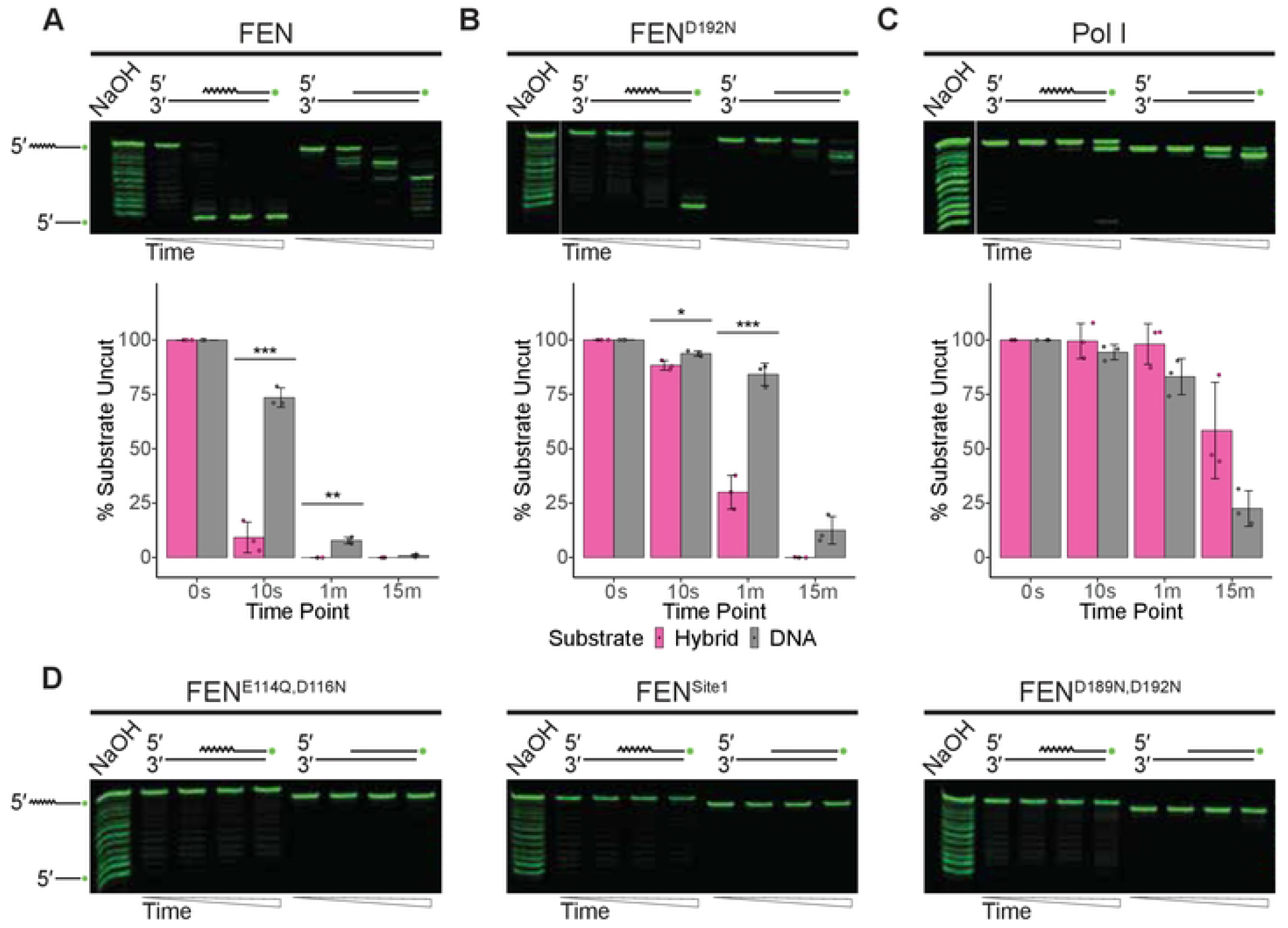
FEN is more active on 3′ overhangs than Pol I. (A) Nuclease activity of FEN, (B) FEN^D192N^, or (C) Pol I on 3′ overhang structures shown using 20% urea-PAGE. Percent of substrate left uncut by the protein, based on three replicates, is shown beneath the respective gel. Significance is indicated by asterisks: * p<0.05, ** p<0.01, or *** p<0.001. Standard error across replicates is indicated by the black bars. (D) Failure of FEN mutants FEN^E1414Q,D116N^, FEN^Site1^, and FEN^D189N,D192N^ to cleave 3′ overhang substrates. Substrates assayed were generated by annealing oJR340 with oJR339 (RNA-DNA hybrid; RNA indicated by zigzags) or oJR348 (DNA). The triangle indicates time point progression of 0 s, 10 s, 1 m, and 15 m; the ladder was produced via alkaline hydrolysis of the hybrid substrate.

### FEN is more active on blunt substrates than Pol I

In addition to Okazaki fragment-like substrates, we assayed the proteins for activity on less canonical substrates. The first of these (**Fig 5**) mimicked blunt DNA, which can occur *in vivo* when a nick is converted to a double stranded break when encountered by a replication fork [59]. As **Fig 5A** shows, FEN had high activity on both the hybrid and DNA-only substrates, with a significant preference for the hybrid. Furthermore, FEN demonstrates different activities on the two variants of the substrate: on the hybrid it showed endonuclease activity and cleaved all the rNMPs while on the DNA-only version it cleaved exonucleolytically. The catalytically reduced FEN^D192N^ mutant produced similar results (**Fig 5B**), although the enzymatic efficiency was reduced. Pol I had no remarkable activity on either substrate (**Fig 5C**). The FEN mutants were not active on the blunt substrates (**Fig 5D**).

**Fig 5.**
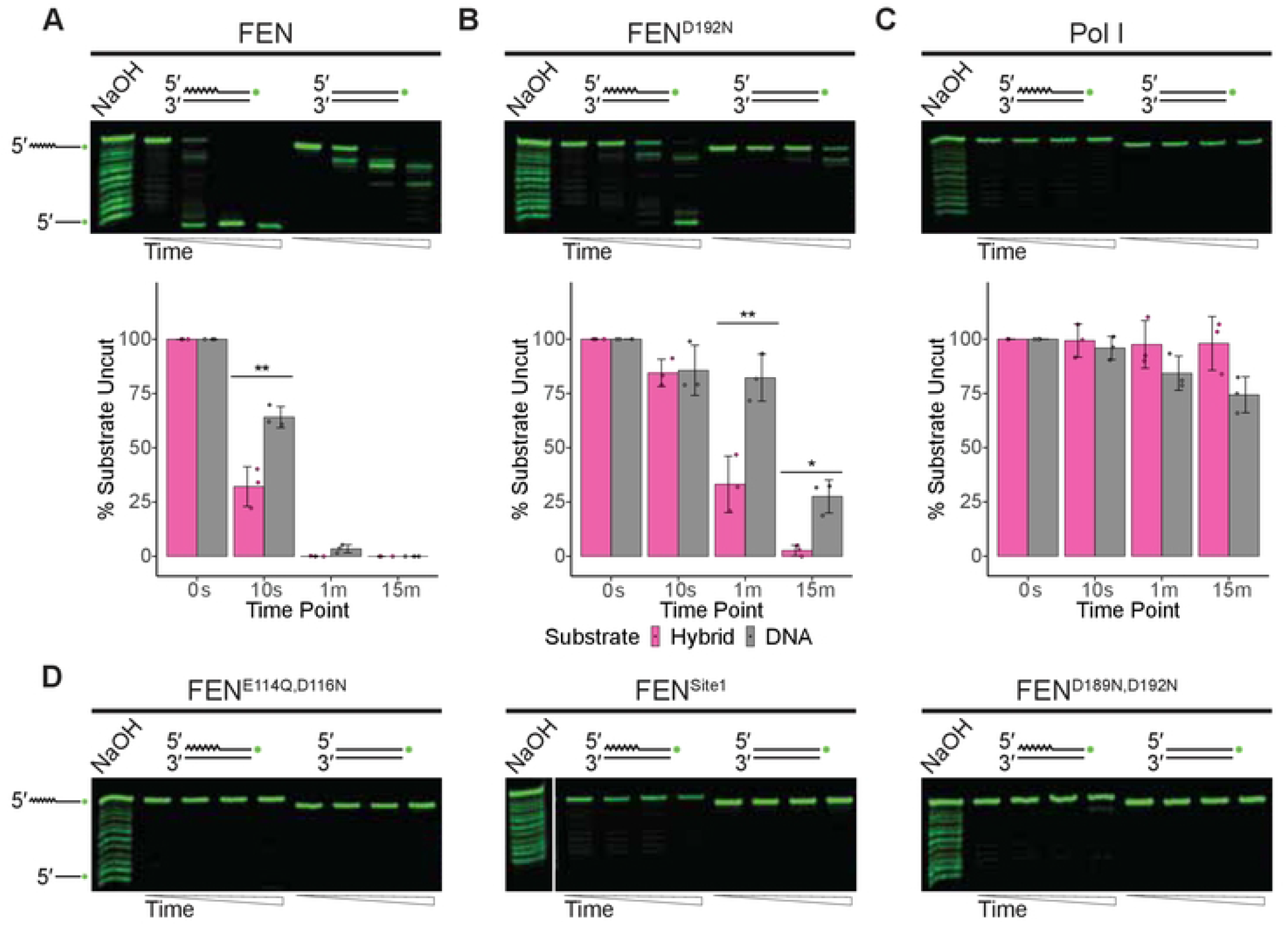
FEN has strong activity on blunt duplex substrates. (A) Denaturing urea-PAGE gels showing assays of FEN, (B) FEN^D192N^, or (C) Pol I activity on blunt duplex structures. Percent uncut substrate remaining from three replicates is quantified under the appropriate gel. Black bars represent standard error and asterisks symbolize significance: * p<0.05, ** p<0.01, or *** p<0.001. (D) Assays of FEN mutants: FEN^E1414Q,D116N^, FEN^Site1^, and FEN^D189N,D192N^ on the blunt substrates. Substrates were composed of oligonucleotides oJR365 and either oJR339 (RNA-DNA hybrid; RNA indicated by zigzags) or oJR348 (DNA only), with a ladder generated by treating the hybrid version with sodium hydroxide. Time points for all reactions are 0 s, 10 s, 1 m, and 15 m.

### Pol I has more activity on 5′ overhangs than FEN

We also tested activity on a 5′ overhang structures (**Fig 6**), which make up a portion of double stranded breaks in the cell [59]. While not as striking as with the previous substrates, FEN showed exonuclease activity on both the hybrid and the DNA-only substrate (**Fig 6A**). This activity was not detected with the FEN^D192N^ mutant (**Fig 6B**), suggesting that the 5′ overhang is not a preferred substrate of FEN. The remaining FEN mutants also had no discernable activity on the substrates (**Fig 6D**). Unlike FEN, Pol I was able to perform endonucleolytic cleavage on both the hybrid structure and the DNA-only 5′ overhang (**Fig 6C**). Pol I had overall lower activity on the DNA-only substrate than the hybrid, however, a distinct band indicative of the removal of multiple rNMPs was present for both. The differences between the activities of FEN and Pol I on the 5′ overhang substrate are striking. Interestingly, FEN from bacteriophage T5 was shown to have activity on 5′ overhangs similar to what is seen here with Pol I, while FEN from *Staphylococcus aureus* behaved similarly to *B. subtilis* FEN [29]. This leads us to conclude that, while FEN and Pol I belong to the same family, these proteins can also have distinct roles in maintaining genome integrity.

**Fig 6.**
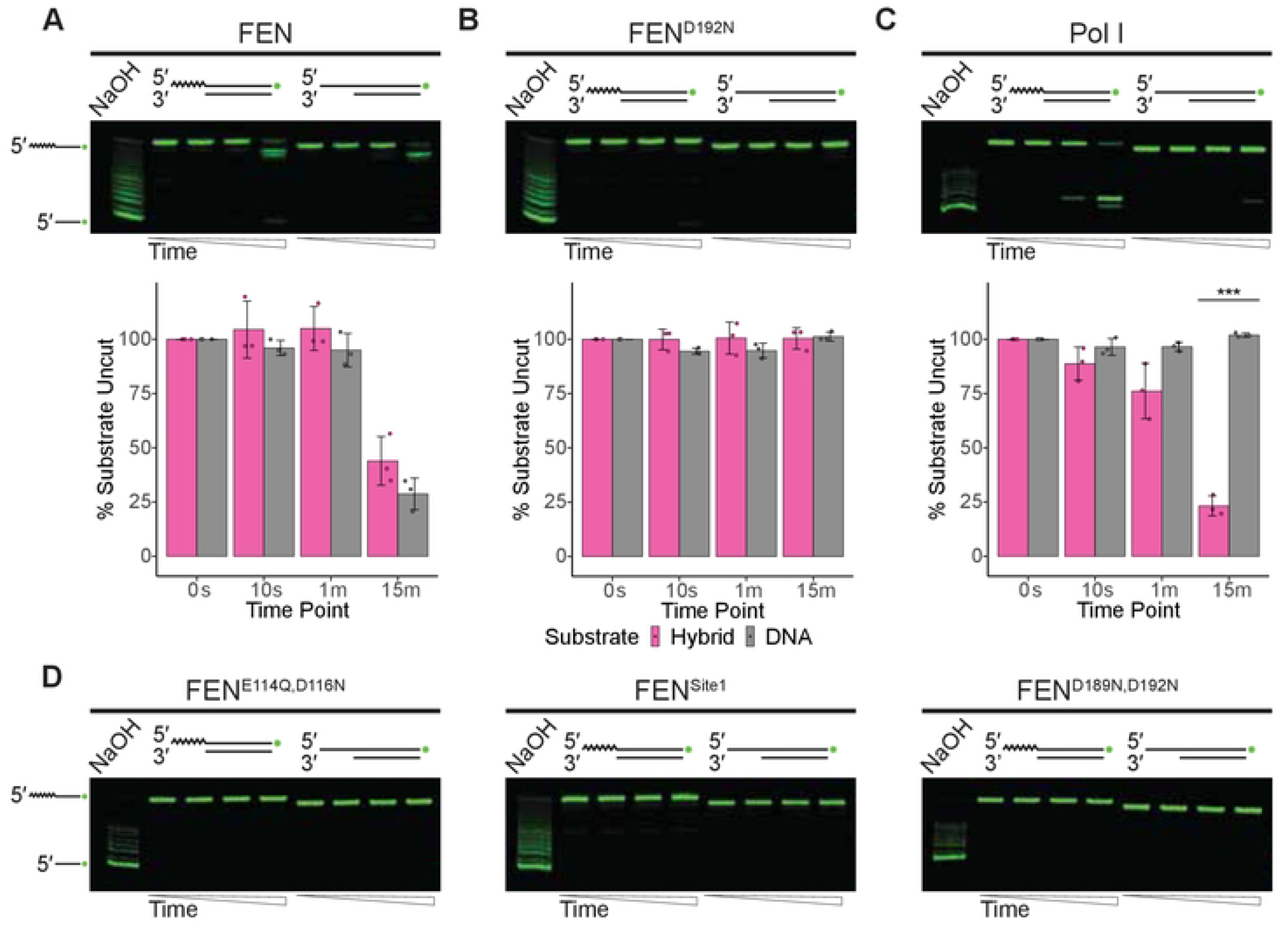
FEN and Pol I have different nuclease activities on 5′ overhang structures. (A) Nuclease activity assays of FEN, (B) FEN^D192N^, or (C) Pol I on duplex DNA with a 5′ overhang resolved using urea-PAGE. Percent substrate left intact by each protein was quantified from three replicates, shown underneath the respective assay. Standard error bars are provided, and statistical significance is indicated by asterisks as follows:: * p<0.05, ** p<0.01, or *** p<0.001. (D) The FEN mutants FEN^E1414Q,D116N^, FEN^Site1^, and FEN^D189N,D192N^ were also assayed for activity on the 5′ overhang structures. 5′ overhang structures were generated by annealing oJR365 with oFCL5 (RNA-DNA hybrid; RNA indicated by zigzags) or oFCL4 (DNA). Reaction time points are 0 s, 10 s, 1 m, and 15 m. Ladder was generated via alkaline hydrolysis of the hybrid 5′ overhang structure.

### FEN has more activity on single-stranded substrates than Pol I

The last construct that we investigated is a single-stranded substrate with no double-stranded regions (**Fig 7**). It has been established that *Ec*Pol I 5′ to 3′ exonuclease activity requires duplex DNA [60], however this is not the case for all FEN proteins [29]. FEN had exonuclease activity on both versions of this substrate, and there was no detectable band indicative of full removal of the rNMPs (**Fig 7A**). The FEN^D192N^ mutant behaved similarly to WT FEN, but again with reduced overall activity (**Fig 7B**). Pol I had negligible activity on either variant and did not create a distinct product, in agreement with prior work showing that neither Pol I from *E. coli* nor *Thermus aquaticus* is active on ssDNA [60,61]. While the three catalytically inactive FEN mutants initially appear to have some activity on this substrate (**Fig 7D and S7A Fig**), assays lacking protein suggest that minor loss of signal is attributable to background degradation of the RNA substrate (**S7B Fig**) as it is more prone to spontaneous hydrolysis when single-stranded. We conclude that FEN is active on ssDNA and ssRNA/DNA hybrids.

**Fig 7.**
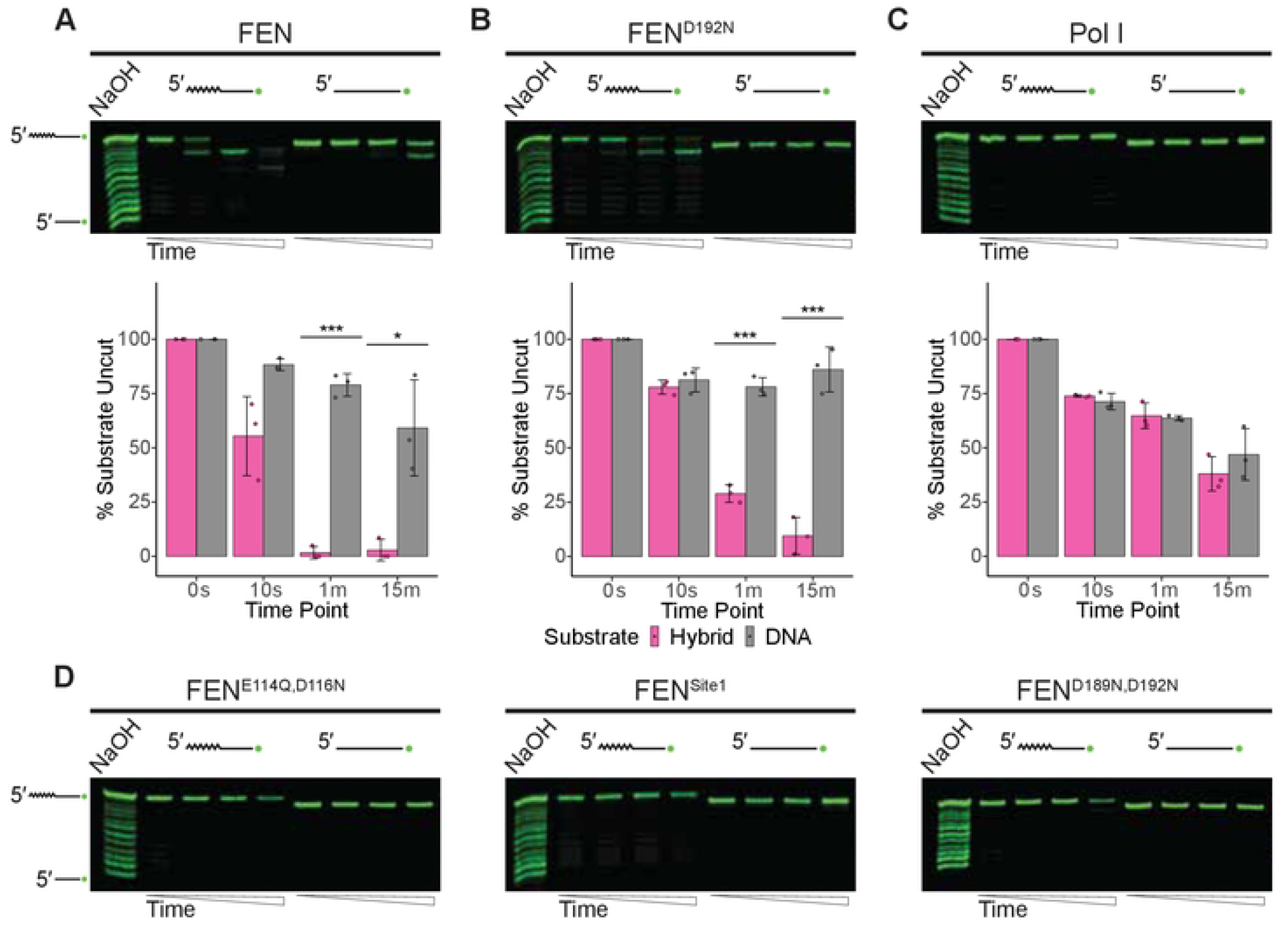
FEN has more nuclease activity on single-stranded substrates than Pol I. (A) Activity of FEN, (B) FEN^D192N^, or (C) Pol I on single-stranded RNA-DNA hybrid or DNA. Percent of intact substrate was quantified from three replicates and shown below the appropriate gel. Significance is indicated by asterisks (* p<0.05, ** p<0.01, or *** p<0.001) and standard error bars are provided. (D) FEN mutants were assayed on the same substrate, with representative gels shown. Single-stranded RNA-DNA hybrid (RNA indicated by zigzags) was oligonucleotide oJR339 while single-stranded DNA was oJR348.

### Pol I nuclease activity is not greatly stimulated with extension

Given the striking differences in the nuclease activities of FEN and Pol I, we asked whether these differences could be attributed to the presence of the Klenow fragment on Pol I. While FEN is composed of a single domain, Pol I also has the Klenow fragment, which is approximately twice as large as the Pol I FEN-domain. In the established model for Okazaki fragment maturation, there must be coordination between the FEN-domain and the Klenow fragment [21,57,62]. To account for this, we tested Pol I activity on a dual-labeled substrate mimicking an Okazaki fragment. As **Fig 8** shows, in the absence of dNTPs, Pol I was unable to extend the free 3′ end of the 5′-labeled substrate and did not noticeably cleave the 3′-labeled substrate. In the presence of dNTPs, Pol I was able to extend the 5’-labeled substrate and cleave some of the downstream nucleotides, although most of the substrate remained intact. We assayed FEN under the same conditions and found that it cleaved the primer-like portion of the 3′-labeled substrate regardless of whether dNTPs were included in the reaction buffer. Thus, even under conditions that promote engagement of both activities of Pol I, FEN had greater catalytic activity on the substrate, which is consistent with our biochemical characterization of FEN. Together, our biochemical work suggests that FEN is the major contributor to the repair of Okazaki fragments in *B. subtilis*.

**Fig 8.**
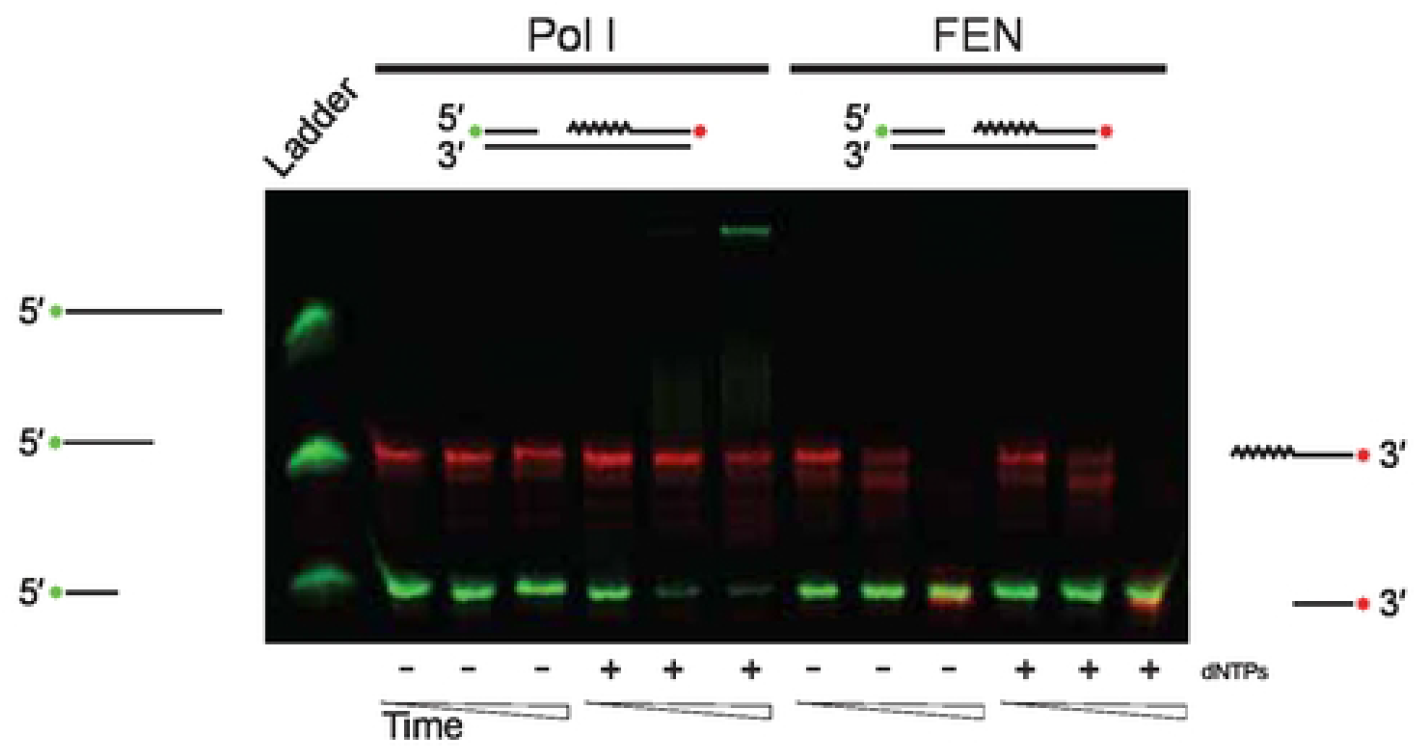
Pol I has lower nuclease activity than FEN even with concurrent DNA synthesis. Denaturing urea-PAGE of products generated by Pol I or FEN activity on a substrate resembling an Okazaki fragment, where upstream and downstream fragments are separated by ten bases. Extension by synthesis from the upstream, 3′ hydroxyl group is tracked by a 5′ fluorescent labeled substrate (green, oFCL8) while nuclease activity on the downstream Okazaki-fragment is tracked with the red colored, 3′ labeled oligonucleotide (oJR367). Labeled oligonucleotides were annealed to oFCL6. Each protein was tested with or without 50 μM dNTPs and time points of 10 s, 1 m, and 15 m. Ladder was generated by adding oJR363 and oJR364 to sodium hydroxide treated substrate.

### Δ*polA* strain sensitivity is due to loss of the Klenow fragment

Unlike the Δ*fenA* strain described earlier, a single deletion of *polA* results in sensitivity to DNA damage from MMC and MMS [13,23,35]. Given our biochemical results, we hypothesized that the Δ*polA* phenotype is due to the loss of the Klenow fragment rather than loss of the FEN-domain. Since prior work has shown that Pol I activity can be reconstituted even with the fragments physically separated [17], we constructed IPTG-inducible strains to express *polA*, the Pol I FEN-domain (*polA*_*FEN*_), or the Pol I Klenow fragment (*polA*_*Klenow*_) in the Δ*polA* background. As shown in **Fig 9**, overexpression of either full-length Pol I or the Klenow fragment was sufficient to rescue the *polA* deletion strain. Overexpressing the Pol I FEN domain did not reduce MMC sensitivity, indicating that the strong phenotypes associated with Δ*polA* strains exposed to DNA damage is due to loss of polymerase activity rather than loss of the protein’s nuclease activity. This result reaffirms that the FEN domain of Pol I is not critical for normal cell growth, and that FEN is sufficient even when cells are exposed to DNA damage.

**Fig 9.**
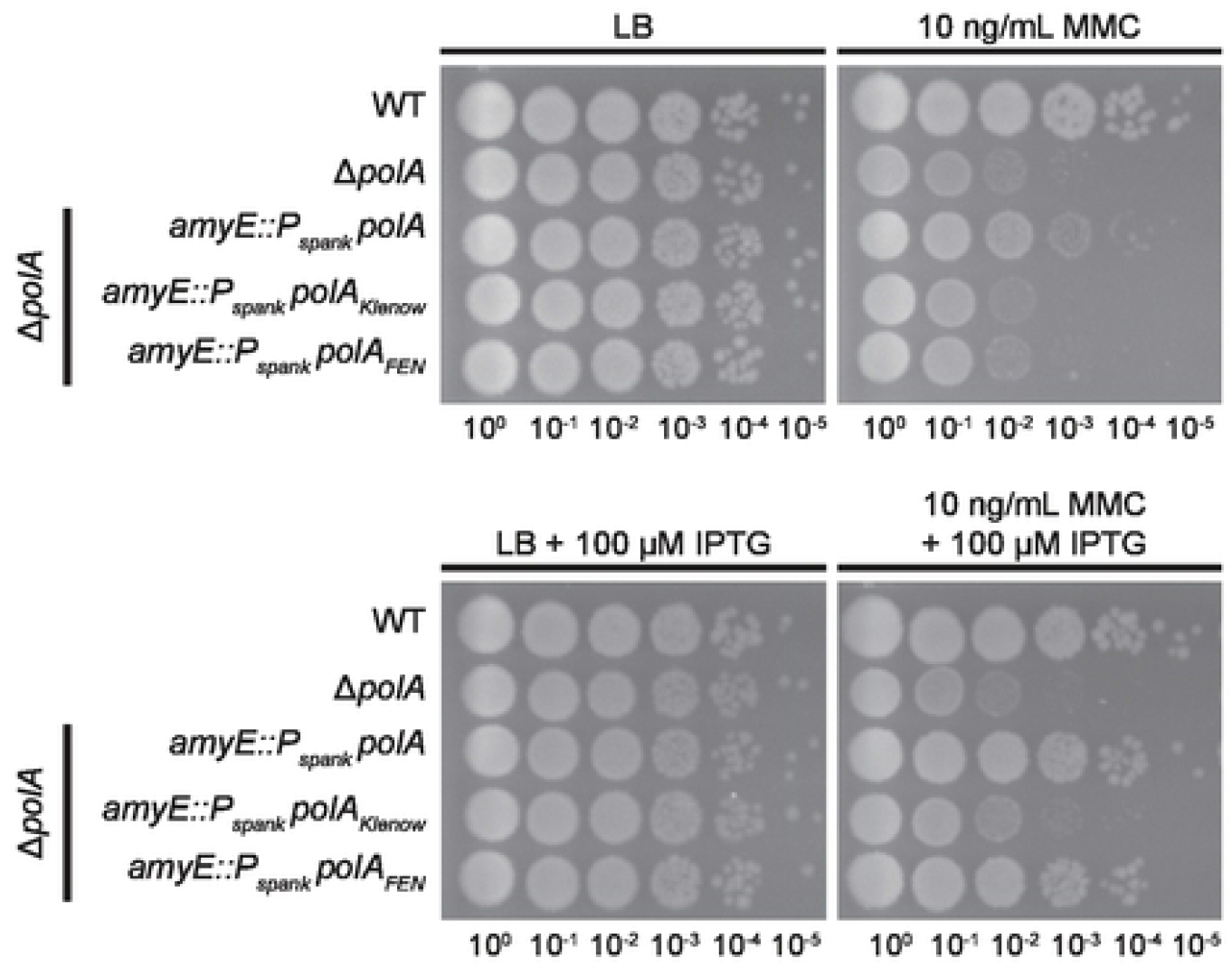
Sensitivity of *polA* deletion strains is due to loss of the Klenow fragment. Rescue of *polA* deletion strain sensitivity to mitomycin C (MMC) by ectopic, full-length Pol I (*polA)* or Pol I polymerase domains (*polA*_*Klenow*_ and *polA*_*FEN*_). Plates were incubated at 30°C.

### Pol I cannot directly substitute for FEN

We have established that, in general, FEN has more nuclease activity than Pol I on biologically relevant substrates assayed *in vitro*. We considered the implications that this might have in the cell. As **Fig 1D** demonstrated, overexpression of FEN or the catalytically compromised FEN^D192N^ mutant was able to rescue the Δ*rnhC, ΔfenA* strain. This led us to question if the biochemical difference we show between FEN and Pol I was realized in cells. We created Δ*rnhC, ΔfenA* strains expressing IPTG-inducible *fenA, fenA*^*D192N*^, *polA, polA*_*FEN*_, or *E. coli xni* and assayed for rescue of HU sensitivity. As before, overexpression of either *fenA* or *fenA*^*D192N*^ resulted in rescue (**Fig 10**). Neither the FEN domain of Pol I nor full-length Pol I were able to rescue the strain as well as *fenA* or *fenA*^*D192N*^, indicating that FEN’s function in cells cannot be fully compensated for by Pol I alone. Expression of *xni*, which encodes the *E. coli* FEN paralog ExoIX, also failed to rescue the strain. This protein has been shown to have some catalytic activity [41] although it is missing three of the carboxylate residues that typify bacterial FENs. (**S2 Fig**); as such, it may have reduced activity compared to FENs that contain all eight residues [29]. We conclude that the high catalytic activity of FEN cannot be directly substituted for by Pol I, especially in the absence of RNase HIII.

**Fig 10.**
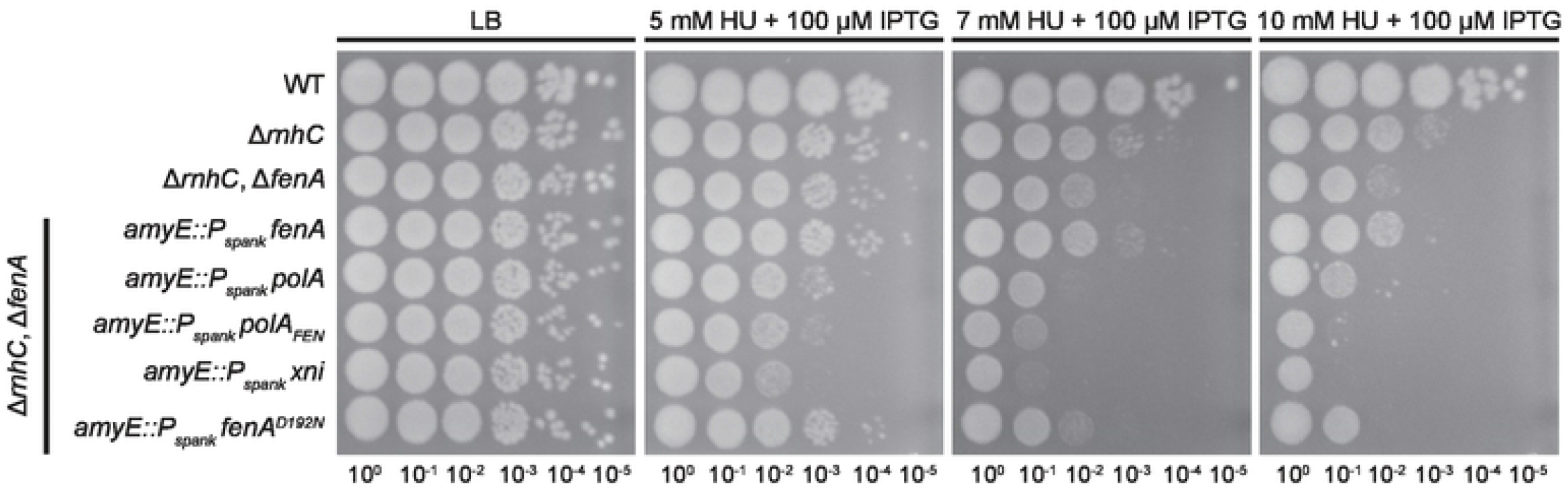
Expression of *polA* does not rescue deletions of *fenA*. Ectopic expression of different proteins with FEN activity in a Δ*rnhC*, Δ*fenA* strain sensitive to hydroxyurea (HU). *fenA* and *polA* encode the full-length proteins from *B. subtilis, fenA*^*D192N*^ and *polA*_*Klenow*_ express derivatives of the native protein, and *xni* encodes the *E. coli* homolog of FEN, ExoIX, which lacks three of the carboxylate residues associated with bacterial FENs.

## Discussion

FEN, previously referred to as YpcP or ExoR, is a member of the large FEN family of proteins that are found across all domains of life [45,63]. These proteins play an essential role in genome maintenance, particularly during Okazaki fragment maturation [64]. For bacteria that encode two proteins with FEN activity, at least one must be present for viability; in *B. subtilis*, this pair is FEN and Pol I [35]. Our biochemical analysis reveals surprising differences between the nuclease activities of FEN and Pol I. We show that, in general, FEN has more nuclease activity than Pol I. This is the case even for canonical structures associated with Pol I activity, such as 5′ flap overhangs and nicks. Across all structures tested, except the 5′ overhang, FEN showed a strong preference for RNA-DNA hybrids over dsDNA. We conclude that FEN’s major biological function is the removal of RNA primers during lagging strand synthesis. There is evidence suggesting that FEN contributes to DNA repair, particularly during UV damage [48,49]. Given the poor activity of FEN on DNA substrates and the lack of a DNA damage phenotype for Δ*fenA*, we conclude that FEN does not provide a significant contribution to the repair of UV-induced DNA damage.

An important consideration regarding the activities of FEN and Pol I is that the latter is physically constrained by the Klenow fragment. The two activities of Pol I are present in separate domains, however they are joined by a flexible linker. Biophysical modeling suggests that the Klenow fragment has lower affinity for the flap structure, which allows it to be replaced by the FEN domain [21,62,65,66]. To account for potential interference by the Klenow fragment, we also assayed for nuclease activity under conditions where extension occurs from an upstream 3′ hydroxyl. While extension by Pol I occurred in the presence of dNTPs, the nuclease activity of Pol I remained lower than that of FEN under the same conditions. The relatively low levels of Pol I nuclease activity that we observed are consistent with other studies, as purifications of Pol I from *B. subtilis* were noted to have lower levels of nuclease activity than purifications from *E. coli* [26,67]. It is important to note that these early results were possibly influenced by proteolysis during protein purification [17,27,68]; to avoid this, the proteins used in this study were confirmed to be full length using SDS-PAGE (**S6 Fig**). Additionally, our group has previously shown that cleavage of an Okazaki fragment-like substrate by Pol I is minimal, and that repair of that substrate is increased when RNase HIII is included in the reaction [13]. Together, our biochemical data show that FEN is the major nuclease during Okazaki fragment maturation in *B. subtilis*, a role previously attributed to DNA polymerase I.

Despite FENs apparent role in Okazaki fragment maturation, Δ*fenA* cells do not demonstrate a detectable phenotype, though a Δ*rnhC*, Δf*enA* strain is more sensitive to genotoxic stress than either of the single deletions. This suggests that FEN is actively involved in the resolution of RNA-DNA hybrids, although some of this activity can be compensated for by other repair pathways [13]. Unlike the Δ*fenA* strain, Δ*polA* strains are sensitized to DNA damage, including damage caused by MMC and UV [13,23,35]. Given our biochemical results, we suspect that this was due to a loss of the polymerization activities of Pol I rather than loss of the nuclease. Indeed, overexpression of the Klenow fragment rescued the Δ*polA* strain as well as full-length *polA*. Similarly, MMS sensitivity in *E. coli* was attributed to mutations affecting the polymerase activity of Pol I [25,69]. Given that Pol I is known to be broadly involved in the repair of DNA damage [55], it is likely that loss of the Klenow fragment and its strand-displacement synthesis activity leads to an accumulation of unrepaired DNA, resulting in genotoxic stress.

It has been accepted that Pol I is the major polymerase involved with Okazaki fragment maturation in bacteria [15,18], due to its dual function as polymerase and nuclease [15]. We show that Δ*rnhC*, Δ*fenA* strains are best rescued by WT FEN or a mutant with reduced catalytic activity (FEN^D192N^), and that overexpression of Pol I or Pol I_FEN_ fails to rescue the strain. Our work complements the results of Fukushima *et. al*, and further advances the idea that, while FEN and Pol I can degrade the same canonical substrates, the proteins have different contributions and substrate preferences in the cell. We suggest that the removal of RNA primers during Okazaki fragment maturation in *B. subtilis* is carried out primarily by FEN while the upstream Okazaki fragment is extended by Pol I, as modeled in **Fig 11**. In the absence of FEN, the combined activities of Pol I and RNHIII could operate as a secondary pathway for maturation of the fragments. We show that Pol I has levels of biochemical activity very similar to FEN^D192N^, yet *fen*^*D192N*^ complements the Δ*rnhC*, Δf*enA* phenotype better than *polA*. We interpret this result to mean that while a complete loss of 5′ nuclease activity from FEN and Pol I is lethal [35], maximal FEN activity is not necessary for normal growth. We conclude that *B. subtilis* cells grow well when some FEN activity is provided, and the existence of two pathways for Okazaki fragment maturation explains the lack of an obvious phenotype for the Δf*enA* strain.

**Fig 11.**
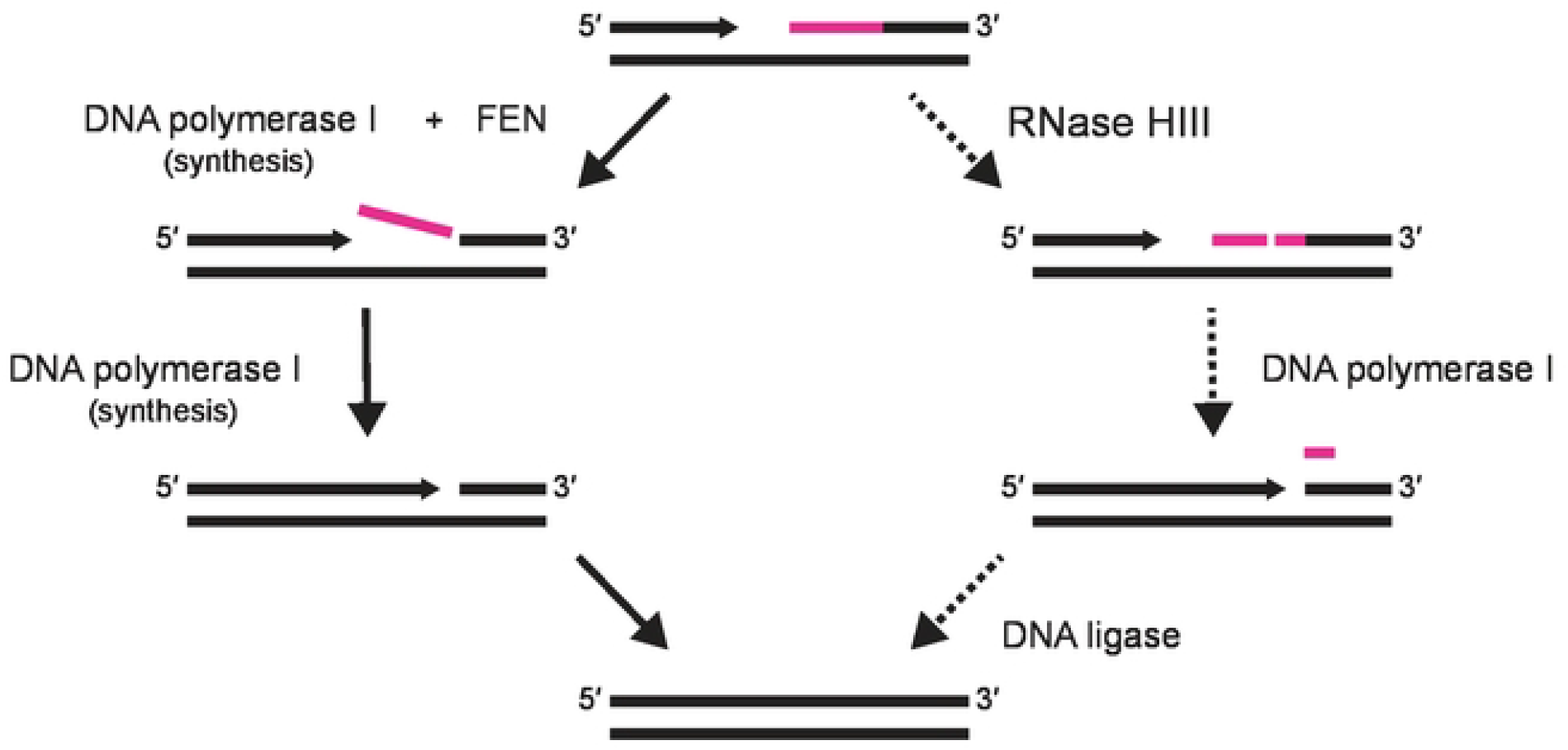
A model of the proposed role of FEN in maturation of Okazaki fragments. During lagging strand synthesis, maturation of Okazaki fragments can occur via two pathways. In the primary pathway (left, solid arrows), FEN removes RNA primers (pink) from Okazaki fragments while Pol I synthesizes from the upstream 3′ hydroxyl. DNA synthesis continues until Pol I loses affinity for the substrate and is outcompeted by DNA ligase, which seals the remaining nick. In the alternate pathway (right, dashed arrows), RNase HIII internally cleaves the RNA primer, shortening the RNA-DNA hybrid to facilitate Pol I nuclease activity. As Pol I approaches the remaining ribonucleotides, it can remove them with its FEN domain following strand-displacement synthesis or nick translation. Synthesis and nucleotide removal continue until Pol I is outcompeted by DNA ligase, which repairs the final nick.

What then is the purpose of *B. subtilis* encoding both Pol I and FEN? Our work shows that the major contribution of Pol I to the cell is through its DNA polymerase domain, for both Okazaki fragment maturation and DNA repair. Based on all available evidence, the contribution of FEN is specific to completion of Okazaki fragments. We propose that the combination of FEN and Pol I provide the most efficient process for primer removal and resynthesis during completion of the lagging strand. This mechanism, with separate proteins contributing FEN and DNA polymerase activity, is reminiscent of the mechanism of Okazaki fragment maturation employed by eukaryotic cells where FEN1 (Rad27) and DNA polymerase δ coordinate during replication [64]. Furthermore, the maintenance of multiple members of the FEN family in the same organism can be seen beyond prokaryotes, as demonstrated by the FEN1 and EXO1 proteins in eukaryotes [70]. As such, the differences between FEN and Pol I that we show here may support a broader pattern of specialized FENs in bacterial DNA replication and repair.

## Methods

### Plasmid Cloning

All primers used are listed in **S2 Table**. Plasmids were constructed using either pDR110 or pE-SUMO as the vector, which were amplified with oJR262 and 263 or oJR46 and 47, respectively. WT *B. subtilis* genes were amplified from PY79 genomic DNA, while FEN mutants were created using site-directed mutagenesis of plasmid carrying the WT gene. *polA*_*FEN*_ or *polA*_*Klenow*_ in the pDR110 vector were generated by selectively amplifying the appropriate regions from the *polA*-expressing plasmid. *xni* was amplified from *E. coli* BL21_DE3_ cells. Genes were inserted into the appropriate vector using Gibson assembly, and subsequently transformed into *E. coli* MC1061 cells. Transformants with spectinomycin (pDR110 vector) or kanamycin (pE-SUMO vector) resistance were confirmed using colony PCR. Plasmid sequences were confirmed via Sanger sequencing.

### *B. subtilis* Strain Construction

All strains used in this work are described in **S3 Table**, and all *B. subtilis* strains are derivatives of PY79. Competent *B. subtilis* was generated by inoculating LB (supplemented with 3 mM Mg_2_SO_4_) with one colony of desired strain and growing in a rolling rack at 37°C until OD_600_ ∼ 0.7. Pre-warmed MD minimal media (1x PC buffer [10x PC buffer: 107 g/L K_2_HPO_4_, 60 g/L KH_2_PO_4_, 10 g/L trisodium citrate • (H_2_O)_5_], 2% glucose, 50 μg/mL tryptophan, 50 μg/mL phenylalanine, 11 μg/mL ferric ammonium citrate, 2.5 μg/mL potassium aspartate, 3 mM MgSO_4_) was inoculated with culture and grown in a rolling rack at 37°C for 6 hours. IPTG-inducible strains were constructed by transforming the appropriate parent strain with the NcoI linearized, pDR110-based plasmid containing the desired gene (plasmids are listed in **S4 Table**). Insertion at the *amyE* locus was confirmed by testing for a loss of starch-hydrolysis and through PCR amplification using oPEB866 and 867.

### Spot Titers

*B. subtilis* strains were streaked onto appropriate LB plates and incubated overnight at 37°C. For each strain, 2 mL of LB was inoculated with a single colony and grown at 37°C in a rolling incubator to OD_600_ 0.5-1.0. Cultures were pelleted at 4000 x g and the supernatant replaced with sterile 0.85% w/v NaCl (saline). Each strain was normalized to OD_600_ = 0.5, then serially diluted to 10^−5^ in sterile saline. 5 μL of each dilution was spotted onto appropriate plates then incubated overnight. Plates were grown at 37°C due to cold sensitivity [13], unless noted to be grown at 30°C. All spot titers were performed in biological triplicate, and all plates were prepared on the day of the experiment. Brightness and contrast adjustment of whole images was performed using ImageJ.

### Protein Expression

Competent BL21_DE3_ (FCL14) were transformed with the appropriate plasmid (**S4 Table**), spread on LB agar supplemented with 25 μg/mL kanamycin, and incubated overnight at 37°C. A starter culture was made by inoculating LB containing 25 μg/mL kanamycin with a single colony, which was grown shaking at 200 rpm in a 37°C incubator overnight. A portion of the starter culture was used to inoculate each liter of LB supplemented with 25 μg/mL kanamycin. Cultures were grown at 37°C with shaking until OD_600_ ∼ 0.6. Each liter of culture expressing FEN or FEN mutants was induced with 0.5 mM IPTG, cooled on ice for 20 min, and grown for 18 hours at 25°C. Cultures expressing Pol I were induced with 0.5 mM IPTG and grown at 37°C for 3 hours. Induced cells were harvested by centrifugation at 4°C and 4000 rpm for 25 min. Supernatant was discarded and pellets were stored at −80°C.

### Protein Purification

Two 1 L pellets were thawed on ice, then resuspended in 60 mL of Lysis Buffer (20 mM Tris, pH 8.0, 400 mM NaCl, 1 mM DTT). One protease inhibitor tablet (Pierce™ Protease Inhibitor Tablets, EDTA-free, A32965) was added, then cells were sonicated on ice for 2.5 minutes total on time (cycles of 10 seconds on and 20 seconds off) at 4°C. Lysed cells were centrifuged at 4°C and 14,000 rpm for 45 minutes, then supernatant was decanted into a clean conical. Ni-NTA agarose (Qiagen, 30210) resin was equilibrated with Lysis Buffer in a gravity column, before clarified lysate was passed over it. The column was washed with 5 volumes of Lysis Buffer, followed by a wash with 10 column volumes of Wash Buffer (20 mM Tris, pH 8.0, 2 M NaCl, 1 mM DTT, 15 mM imidazole). Fractions containing protein were eluted with 3 column volumes of Elution Buffer (20 mM Tris, pH 8.0, 400 mM NaCl, 1 mM DTT, 300 mM imidazole), and the most concentrated fractions were determined using a NanodropLite spectrophotometer. Selected fractions were pooled and treated with SUMO-protease for 2 hours at room temperature to remove the 6x-His-SUMO tag. Digested protein was transferred to 10K molecular cut off weight (MCOW) dialysis tubing and dialyzed overnight against Dialysis Buffer (20 mM Tris, pH 8.0, 300 mM NaCl, 1 mM DTT) at 4°C overnight. The protein was passed over a Ni-NTA gravity column equilibrated with Dialysis Buffer, and the flowthrough was collected. The column was further washed with 3 column volumes each of Dialysis Buffer, Wash Buffer, and Elution Buffer. Each flowthrough was collected, and samples were analyzed using a 10% SDS-PAGE gel stained with Coomassie brilliant blue. For all proteins, the initial flowthrough and dialysis-wash flowthrough was pooled and concentrated using a 10K MCOW centrifugal filter unit (Amicon, UFC9010). Protein was stored in 25% glycerol at −80°C. Further purification was completed using a HiTrap Q FF anion exchange column (Cytivia, 17515601) on an AKTA FPLC using Q Buffer A (20mM Tris, 1mM DTT, 5% glycerol) and Q Buffer B (20mM Tris, 500mM NaCl, 1mM DTT, 5% glycerol). Briefly, the column was equilibrated with 10% Q Buffer B and protein was diluted to 50 mM NaCl in Q Buffer A before loading. Protein was fractionated using an increasing gradient of Q Buffer B. Peak fractions were determined using SDS-PAGE, then peak fractions were pooled and concentrated using a centrifugal filter. Glycerol was added to 25%, followed by flash freezing aliquots in liquid nitrogen before storing at −80°C. Gel demonstrating protein purity was generated using by loading 1 μg of each protein onto a Mini-PROTEAN TGX 4-20% gradient gel (BIO-RAD, 4561096) and then staining with Coomassie blue.

### Nuclease Activity Assays

Substrates were generated by combining 1 μM labeled oligonucleotide with 2 μM of each appropriate unlabeled oligonucleotide in 1x Dilution Buffer (20 mM Tris, pH 8.0, 50 mM NaCl) and boiling for 1 minute at 98°C before being allowed to cool to room temperature. The following oligonucleotides (synthesized by IDT) were used: flap substrates included oJR366, oJR368, and either oJR339 or oJR348, nicked substrates contained oJR338, oJR340 and either oJR339 or oJR348, 3′ overhang structures consisted of oJR340 and either oJR339 or oJR348, blunt substrates consisted of oJR365 and either oJR339 or oJR348, 5′ overhang structures were oJR365 with oFCL4 or oFCL5, and single-stranded oligos were either oJR339 or oJR348. Sequences for all oligonucleotides are listed in **S1 Table**. Purified proteins were diluted to 250 nM in 1x Metals Buffer (20 mM Tris, pH 8.0, 50 nM NaCl, 1 mM MgCl_2_, and 10 μM MnCl_2_). Each reaction contained 100 nM substrate, 50 nM protein and 1x Metals Buffer. Assays were performed at 25°C, and samples were added to an equivalent volume of Stop Buffer (95% formamide, 20nM EDTA, bromophenol blue) at 0 s, 10 s, 1 min, and 15 min. Stopped samples were incubated at 98°C for 5 minutes, immediately followed by snap-cooling on ice. Each ladder was generated by incubating 500 nM of the chimeric form of a substrate in 200 nM NaOH at 37°C for 8 minutes (5 minutes for the 5′ overhang), then stopped as described. Products were resolved using 20% denaturing urea-PAGE and visualized using the 800 nM channel of a LiCor Odyssey imager. Each reaction was repeated at least three times, using different preparations of substrate and different aliquots of protein.

### Analysis of Nuclease Activity Assays

After a gel image was opened in ImageJ, the 0 s lane was selected and identified as the reference lane. The 10 s, 1 min, and 15 min lanes were also selected, and the lane intensities were plotted. Due to manufacturing limitations, the hybrid structure produces more background noise than the DNA-only substrates. The area under each curve was recorded using the Wand Tool, then measurements for each combination of protein/substrate/type were collected from three replicates and exported. Relative percent of uncut substrate was calculated by dividing the area for each timepoint by the area of the corresponding 0 s timepoint. Plots were made using ggplot2 [71] in R [72]; differences in protein activity on RNA-DNA hybrid substrate and DNA-only substrate were determined using Welch’s T Test for unequal variance, an alpha of 0.05, and n=3.

### Okazaki Repair Assays

Repair assays were carried out using the method previously described [12,13]. Briefly, substrates were generated by combining 1 μM oJR367, 1 μM oFCL8, and 2 μM oFCL6 in 1x Dilution Buffer then boiling for 1 minute at 98°C and cooling to room temperature [9,13]. As noted in **S1 Table**, oFCL8 contains phosphorthioate bonds at the 5’ end to prevent off-target 5’ exonuclease activity. Reactions were carried out in 1x Extension Buffer (40 mM Tris-acetate, pH 7.8, 12 mM magnesium acetate, 300 mM potassium glutamate, 3μM ZnSO4, 2% (w/v) polyethylene glycol, 0.02% pluronic F68), using 50 nM protein and 100 nM substrate, with or without 50 μM deoxynucleotide triphosphates (dNTPs). Assays were completed as described above for the Nuclease Activity Assays; however, an appropriate ladder was generated by combining the substrate (treated with sodium hydroxide) with an equal volume of 0.5 μM oJR363 and oJR364 annealed to oFCL6. The dual-colored substrate was imaged using the 700 nM and 800 nM channels of a LiCor Odyssey imager.

## Acknowledgements

We would like to thank members of the Simmons lab for helpful discussions during the progression of this work.

## Supporting Information

**S1 Fig. Strains with single deletions of *fenA* are not sensitive to UV damage**. Spot titer assay of strains exposed to UV damage. RNase HIII (*rnhC*) is a protein known to repair RNA-DNA hybrids in *B. subtilis*.

**S2 Fig. The sequence of FEN contains eight carboxylate residues associated with bacterial FENs**. Multiple sequence alignment of Pol I (N-terminal FEN domain) and FEN from *B. subtilis* as well as the FEN homolog from *E. coli*, ExoIX. Conserved residues are boxed in grey while the active site carboxylate residues that coordinate metal-binding are boxed in pink.

**S3 Fig. FEN activity is dependent on conserved carboxylate residues**. Ectopic expression of *fenA* and *fenA* mutants in the hydroxyurea-sensitive Δ*rnhC*, Δ*fenA* strain.

**S4 Fig. Dominant negative phenotype of *fenA***^***E114Q***,***D116N***^ **and *fenA***^***D189N***,***D192N***^ **is not detectable in a WT background**. WT *B. subtilis* ectopically expressing *fenA* or *fenA* mutants with the indicated changes. Cells were imaged after growth at 30°C for 16 hours.

**S5 Fig. *fenA***^***E114Q***,***D116N***^ **and *fenA***^***D189N***,***D192N***^ **are dominant negative in the absence of *rnhC***. Overexpression of ectopic *fenA* or *fenA* mutants in cells lacking *rnhC*.

**S6 Fig. Proteins used in assays were the major product of relative purifications**. 1 μg of each protein for *in vitro* assays purified as described in Methods section. SDS-PAGE gels were visualized with Coomassie brilliant blue.

**S7 Fig. Activity of FEN mutants on single-stranded substrates is similar to spontaneous substrate degradation**. (A) Quantification of three replicates of nuclease assays using FEN^E1414Q,D116N^, FEN^Site1^, or FEN^D189N,D192N^. Statistical significance is indicated by asterisks corresponding to p<0.05 (*), p<0.01 (**), or p<0.001 (***) with standard error bars provided. (B) Activity assay was repeated without the addition of protein (left) and percent substrate remaining intact was visualized graphically (right). Substrates are oJR339 (RNA-DNA hybrid; RNA indicated by zigzags) and oJR348 (DNA).

**S1 Table. Oligonucleotides used in *in vitro* assays**.

**S2 Table. All primers used in this study**.

**S3 Table. All strains used in this study**.

**S4 Table. All plasmids used in this study**.

